# *Shigella* serotypes associated with carriage in humans establish persistent infection in zebrafish

**DOI:** 10.1101/2022.11.08.515620

**Authors:** Vincenzo Torraca, Dominik Brokatzky, Sydney L. Miles, Charlotte E. Chong, P. Malaka De Silva, Stephen Baker, Claire Jenkins, Kathryn E. Holt, Kate S. Baker, Serge Mostowy

**Author notes:** **Correspondence** (V.T.), (S.M.).

## Abstract

*Shigella* represents a paraphyletic group of human-adapted *Escherichia coli* lineages that converged towards the same enteropathogenic pathovar. In low-income countries, *Shigella* is typically transmitted by ingestion of contaminated water and food, while in high-income countries sexual transmission among men who have sex with men (MSM) is the dominant route of transmission. Although more than 40 different serotypes of *Shigella* have been reported globally, within the MSM community the majority of reported cases are attributed to three *Shigella* serotypes: the *Shigella sonnei* unique serotype (Ss) and the *Shigella flexneri* serotypes 2a and 3a. Here, using a zebrafish infection model, we demonstrate that *Shigella* can establish persistent infection *in vivo*. In this case, bacteria are not cleared by the host immune system and become tolerant to therapeutic doses of antibiotics. We show that persistence is dependent on the *Shigella* O-Antigen, a key constituent of the bacterial cell surface and determinant of serotype classification. Representative isolates of three *Shigella* serotypes associated with global dissemination and MSM transmission (*S. sonnei* Ss, *S. flexneri* 2a and 3a) all persist in zebrafish, while a serotype not associated with MSM transmission (*S. flexneri* 5a) does not. Strikingly, *Shigella* serotypes which establish persistent infection fail to promote macrophage cell death *in vivo*. We conclude that zebrafish can be a valuable platform to illuminate host and pathogen factors underlying the establishment of persistent infection with *Shigella* in humans.

**Highlights:**

- *Shigella* can establish persistent infection in zebrafish
- Establishment of persistent infection *in vivo* depends on *Shigella* serotype
- The same serotypes establishing persistent infection in zebrafish are prevalent in a patient group in whom persistent infection is observed
- *Shigella* serotypes which establish persistent infection fail to promote macrophage cell death *in vivo*

**In Brief:** In high-income countries, sexual transmission among men who have sex with men (MSM) is the dominant cause of domestic *Shigella* transmission. Torraca *et al*. discover that three serotypes of *Shigella* associated with MSM transmission also persist in zebrafish. These results indicate that *Shigella* serotypes associated with MSM transmission can establish carriage in the host, and highlight the use of zebrafish infection to study *Shigella* persistent infection in humans.

## Introduction

Shigellosis is a diarrhoeal disease caused mainly by *Shigella sonnei* and *Shigella flexneri*^1^. Infection from *Shigella* is viewed to be self-limiting and short-lived. In high-income countries, sexual transmission of shigellosis in men who have sex with men (MSM) is the major route of domestic dissemination. In a recent retrospective cohort study performed in the United Kingdom, *Shigella* was longitudinally sampled from patients (up to 176 days apart). In a subset of patients, a small single nucleotide polymorphism (SNP) distance (<8 nucleotides apart) between serial isolates was determined^2^, which is suggestive of persistent carriage. Symptomatic and asymptomatic *Shigella* carriage, as well as prolonged bacterial shedding, have been indicated by culture and molecular testing methods in different populations (Latin America, Sub-Saharan Africa, South Asia and Oceania) and age categories^3–5^. Pathogens recognised to establish persistent infection represent a major public health burden; prominent examples include *Helicobacter pylori, Salmonella enterica* Typhimurium and *Mycobacterium tuberculosis*^6–9^. Persistent infections fail to be cleared by the immune system and are also notoriously recalcitrant to antimicrobial therapy^10^. These infections can become asymptomatic, and together with reactivation at later times (due to a variety of factors, including ageing, immunosuppression and superinfections), can promote future dissemination of the disease.

The zebrafish (*Danio rerio*) is a widely adopted vertebrate model, and valuable to investigate infection by a variety of human bacterial pathogens including enterobacteria, such as *Escherichia coli, Salmonella* and *Shigella*^11,12^. From the very early developmental stages, zebrafish larvae have an innate immune system highly homologous to that of humans, and transparent larvae enable high-resolution intravital imaging of fluorescently-tagged immune cells and bacteria *in vivo*^11,13^. Here, using a *Shigella*-zebrafish infection model, we discover that clinical isolates of *Shigella* can establish a persistent and antibiotic-tolerant infection in zebrafish for at least 6 days (>144 hours). Our results demonstrate a new role for *Shigella* O-Antigen (O-Ag) variants in enabling persistence, and highlight zebrafish as an animal model that can be used to investigate the surge of three dominant *Shigella* serotypes currently circulating in the MSM community^14–16^.

## Results

### *Shigella* can establish persistent infection in zebrafish

To test if *Shigella* can establish persistent infection in zebrafish, we injected a low dose (∼1000 CFU, characteristically eliciting ∼20% host death by 72 hpi) of mCherry-labelled *S. sonnei* 53G in the hindbrain ventricle (HBV) of zebrafish larvae at 3 days post-fertilisation (dpf) and assessed bacterial burden and host survival daily up to 144 hours post-injection (hpi) (**Figure 1A-C, Figure S1A-C**). Quantification of bacterial burden by CFU enumeration (**Figure 1B**) and linear regression analysis (**Figure S1A**) indicate that infection progresses in three distinct phases: an acute phase characterised by bacterial replication (0-24 hpi), a clearance phase characterised by a significant decrease of bacteria at a constant rate (24-96 hpi), and a persistence phase where few bacteria (i.e., <5% of the initial bacterial load) are maintained over an extended period of time (96-144 hpi).

**Figure 1.**
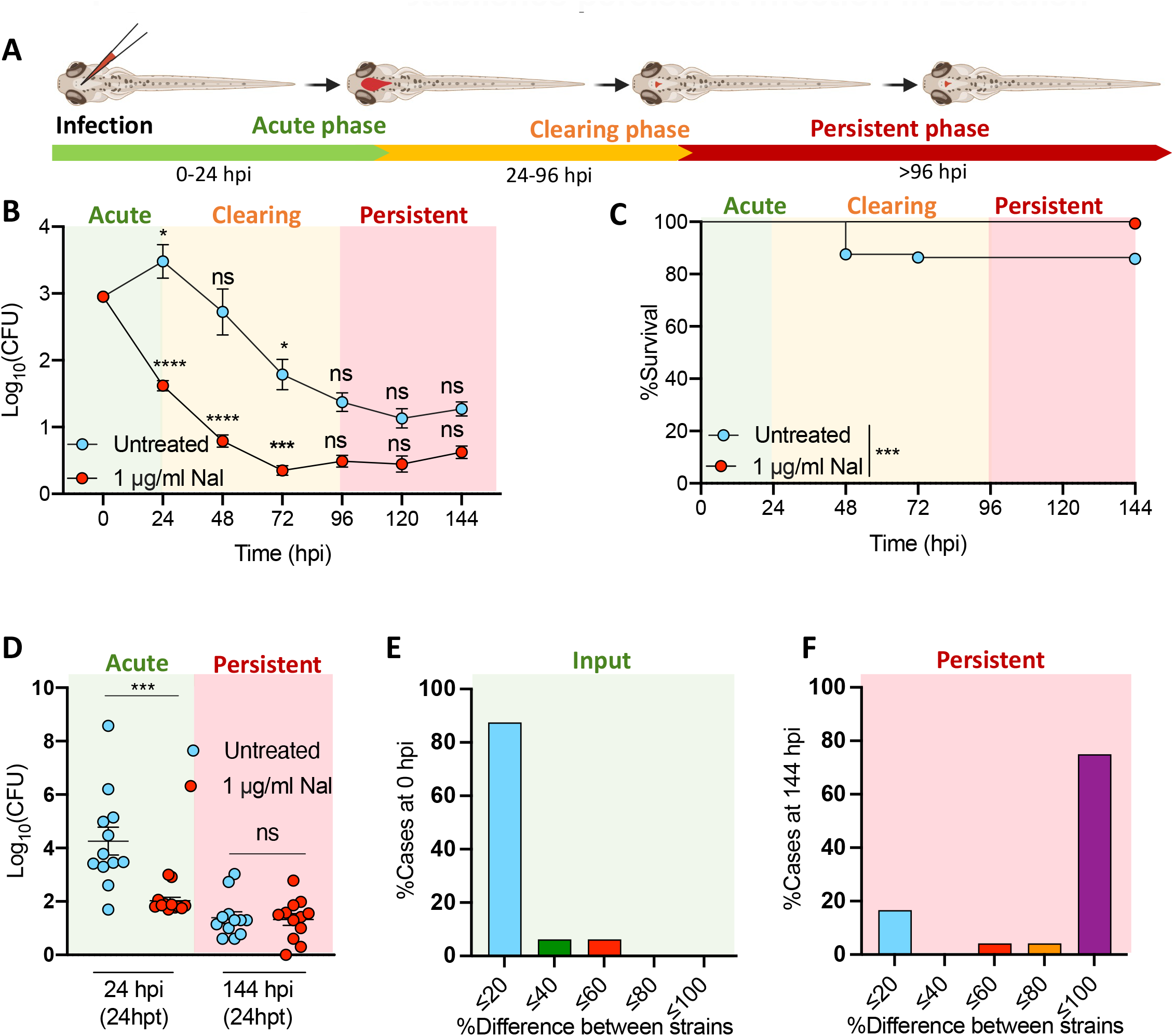
*Shigella* establishes persistent infection in zebrafish. (**A**) Experimental diagram of *S. sonnei*-zebrafish infection. *S. sonnei* infection undergoes three phases, an acute phase (i.e., increasing bacterial load) a clearing phase (i.e., a steady decrease of bacterial load) and a persistent phase (i.e., bacterial load does not further decrease). (**B-C**) CFU count and survival analysis from zebrafish larvae infected with *S. sonnei*. Larvae were treated with the antibiotic Nalidixic acid (Nal) or were left untreated. *S. sonnei* establishes a persistent infection in zebrafish, even during treatment with a therapeutic antibiotic dose. From 96 hours post-infection (hpi), the bacterial load in larvae remains constant over consecutive days. (**D**) CFU count from zebrafish larvae treated with Nal (or left untreated), during acute and persistent infection stages. During the acute infection stage (0 to 24 hpi) *S. sonnei* is sensitive to the antibiotic treatment, while during the persistent infection stages (120 to 144 hpi) *S. sonnei* becomes insensitive to antibiotic treatment. Data were analysed at 24 hours post-treatment (hpt). (**E-F**) mCherry and GFP labelled-*S. sonnei* were co-injected at a 1:1 ratio. At 144 hpi one of the two strains could be isolated at a much higher frequency than the other strain, indicating establishment of clonality. Statistics: unpaired t test on Log10 transformed data against the previous timepoint (B); Log-rank Mantel-Cox test (C); one-way ANOVA with Sidak’s correction (D); ns (non-significant) p≥0.05; *p<0.05; ***p<0.001.

Treatment of infection with a therapeutic dose of the antibiotic Nalidixic acid prevented host death in response to both low dose (**Figure 1B,C)** and high dose (∼8000 CFU, characteristically eliciting ∼80% host death by 72 hpi, **Figure S1D,E**) infections. Treatment with Nalidixic acid also inhibited bacterial growth (**Figure 1B, Figure S1E)** and therefore larvae did not experience an acute phase of infection (as compared to untreated larvae); instead, they immediately progressed towards a clearance phase (**Figure 1B, Figure S1A-C**). In the presence of Nalidixic acid, the rate of bacterial clearance is not significantly different from that of untreated larvae (**Figure S1A-C**), where infection is not fully cleared and has progressed towards a persistence phase. To investigate if bacteria infecting zebrafish in the persistence phase had developed antibiotic tolerance (irrespective of previously being treated with an antibiotic), we treated larvae exhibiting acute and persistent *Shigella* infection with Nalidixic acid for 24h and found that treatment significantly decreased bacterial burden (∼100 fold) during acute infection (**Figure 1D**). In contrast, the bacterial burden in larvae in the persistence phase did not significantly decrease with antibiotic treatment (**Figure 1D**). We observed that spontaneous evolution of antibiotic resistance occurred in 0.53% of larvae (at 72 hpi) undergoing antibiotic treatment, while it was not observed in untreated larvae (**Figure S1G**). This indicates that failure of Nalidixic acid to clear persistent bacteria in most larvae (99.47%) results from antibiotic tolerance.

To study whether persistent bacteria represent a clonally expanded population or derive from a stochastic reduction of the bacterial load, we co-injected *S. sonnei* 53G labelled with either GFP or mCherry (and otherwise isogenic) at a 1:1 ratio (**Figure 1E-F**). As expected, at 0 hpi both strains could be recovered from most larvae (∼90%) at similar loads (difference in the recovery of the two strains ≤ 20%). Strikingly, at 144 hpi, most larvae (∼80%) dominantly carried only one of the two strains (difference in the recovery of the two strains > 80%). These results indicate that persistent *Shigella* infections represent a clonal population.

Together, these data show that *Shigella* can establish persistent infection *in vivo*, which is characterised by poor host clearance over an extended period, antibiotic tolerance and clonality.

### *Shigella* O-Antigen variants circulating in the MSM population promote persistence *in vivo*

To investigate bacterial factors required for persistence *in vivo*, we performed infections using various *S. sonnei* mutants including a Type III Secretion System (T3SS) deficient strain (ΔMxiD), an O-Ag deficient strain (ΔO-Ag) and a strain having lost the virulence plasmid (-pSS). We observed that the ΔMxiD strain can establish persistent infection *in vivo*, however, the ΔO-Ag and -pSS strains cannot (**Figure 2A-B, Figure S2A-B**). Considering that the O-Ag of *S. sonnei* is encoded by the pSS plasmid (i.e., -pSS strains are also O-Ag-deficient), we conclude that the O-Ag (and not the T3SS) is essential for *Shigella* persistence.

**Figure 2.**
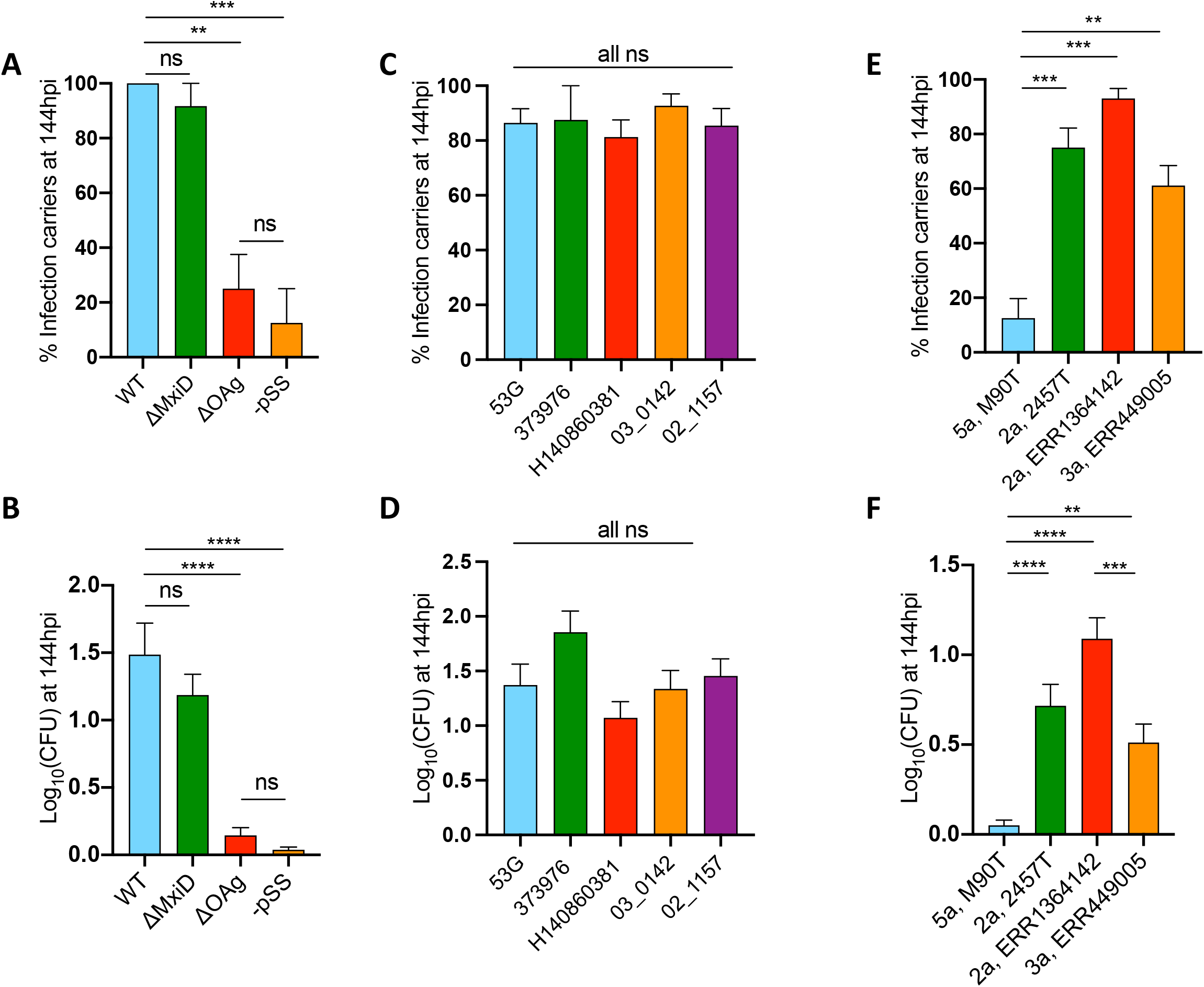
*Shigella* O-Antigen serotypes associated with MSM transmission enable persistent infection. (**A-B**) Percentage of larvae carrying persistent infection at 144 hpi (A) and average bacterial Log_10_CFU (B), for WT *S. sonnei* 53G and several isogenic mutants. A T3SS mutant (ΔMxiD) can establish persistent infection at similar levels as WT. However, the -pSS mutant (depleted of the virulence plasmid) or an O-Antigen mutant (ΔOAg) are significantly reduced in their capability to establish persistent infections. (**C-D**) Percentage of larvae carrying persistent infection at 144 hpi (C) and average bacterial Log_10_CFU (D) for several *S. sonnei* isolates from Lineage II and Lineage III. Both Lineage II and III *S. sonnei* isolates are capable of establishing persistent infections. Although belonging to different lineages, all *S. sonnei* isolates share an identical O-Antigen. (**E-F**) Percentage of larvae carrying persistent infection at 144 hpi (E) and average bacterial Log_10_CFU (F) for several *S. flexneri* isolates from serotypes 2a, 3a and 5a. *S. flexneri* isolates of serotypes 2a and 3a are both capable of establishing persistent infection, to a much greater extent than *S. flexneri* of serotype 5a. Statistics: one-way ANOVA with Sidak’s correction on percentage data (A,C,E) or Log10-transformed data (B, D, F); ns (non-significant) p≥0.05; **p<0.01; ***p<0.001; ****p<0.0001.

More than 40 different serotypes of *Shigella* have been reported, and the O-Ag structure is a major variable in determining these different serotypes^17^. Only three serotypes have been directly associated with carriage in the MSM population to-date^2,18^. To test whether strain clusters associated with carriage in humans establish persistence in zebrafish, we constructed phylogenetic trees for *S. sonnei* and *S. flexneri* using high quality complete genomes deposited in the Bacterial and Viral Bioinformatics Resource Centre (BV-BRC, https://www.bv-brc.org/), sequencing data (short-read assemblies) collected from cases of likely carriage^18^, as well as newly collected sequencing data (long-read assemblies) from *S. sonnei* and *S. flexneri* clinical isolates selected for further testing in zebrafish (**Figure S2G-H**). For the construction of trees, we used the Codon Tree method and the Phylogenetic tree building service made available by BV-BRC^19^. Based on this phylogenetic analysis, we selected *Shigella* isolates that represented either a persistence^20^ ingroup (i.e., falling in a genetic cluster that encompasses isolates previously associated with carriage in humans^18^) or a persistence outgroup (i.e., not directly clustering with isolates previously associated with carriage in humans) for zebrafish infection.

For *S. sonnei*, different sub-lineages have been identified^20–22^, however only one *S. sonnei* serotype exists (**Figure S2G**), which has comparable genetic diversity to individual serotypes of *S. flexneri* (e.g. *S. flexneri* 2a)^23,24^. Both representatives of the persistence ingroup and the persistence outgroup of *S. sonnei* persist in zebrafish *in vivo*. This finding is not surprising, considering that *S. sonnei* isolates have short genetic distances and all share an identical O-Ag that defines the sole *S. sonnei* serotype (**Figure 2C-D, Figure S2C-D**). In the case of *S. flexneri*, multiple serotypes have been identified^25^, although only 2 serotypes (2a and 3a) have been so far associated with persistent carriage^2^ (**Figure S2H**). Strikingly, representative strains of these serotypes also persist in zebrafish (**Figure 2E-F, Figure S2E-F**). In contrast, *S. flexneri* M90T (an isolate belonging to the 5a serotype and a persistence outgroup) is unable to persist in zebrafish (**Figure 2E-F, Figure S2E-F**). Overall, these results indicate a central role for the O-Ag in establishment of persistence. Our findings are consistent with epidemiological evidence showing that transmission in the MSM community is associated with *S. sonnei, S. flexneri 2a*, and *S. flexneri 3a* serotypes, but not *S. flexneri* 5a^2^.

### *Shigella* can establish persistent infection in macrophages *in vivo*

Macrophages are widely viewed as a first line of host defence against *Shigella* infection and have been reported to act as a long-term reservoir for intracellular bacterial pathogens, such as *Salmonella enterica* and *M. tuberculosis*^8,9,26^. Having established a zebrafish model of *Shigella* persistent infection, we sought to determine whether this was localised to macrophages. Using the *Tg(mpeg1::Gal4-FF)*^gl25^/*Tg(UAS:LIFEACT-GFP)*^mu271^ transgenic zebrafish line (labelling macrophages) and mCherry-labelled *S. sonnei* 53G, we identified ∼40% of bacterial fluorescence co-localising with macrophages at 144 hpi (**Figure 3A-B, Video S1**). Longitudinal studies using high-resolution confocal microscopy showed that macrophages harbouring a stable bacterial load (i.e., bacterial fluorescence shows only a 1.21 ± 0.21 fold change over a 12 h period of observation) can be detected as early as 24 hpi (**Figure 3C,Figure S3C, Video S2**) and lasted for several consecutive days (**Figure S3A-B**). These data were surprising, considering that macrophage cell death is widely recognised as a hallmark of *Shigella* infection^12,27,28^.

**Figure 3.**
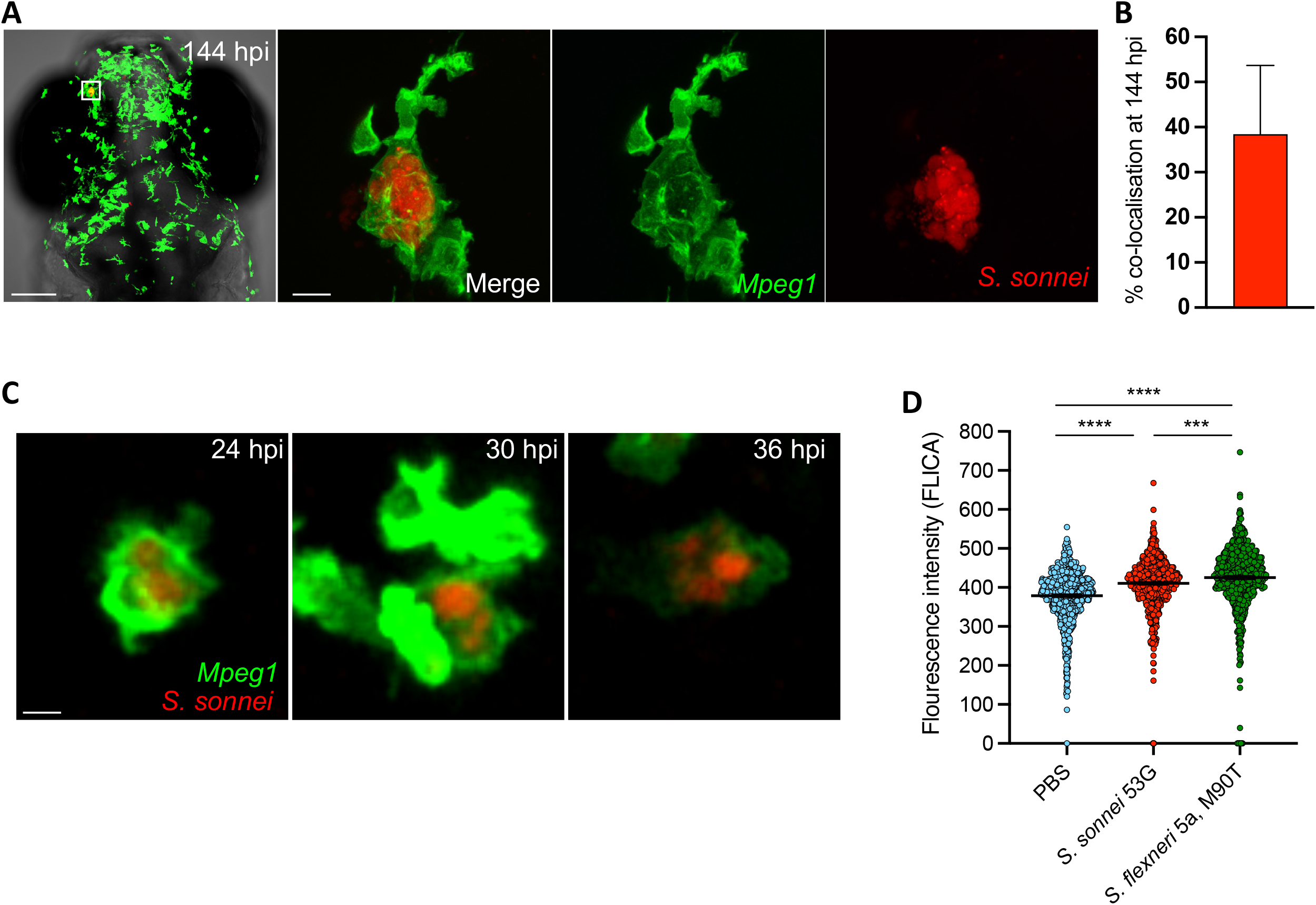
*Shigella* can establish persistent infection of macrophages *in vivo*. (**A**) Head region and individual macrophage detail from a representative *S. sonnei* infected zebrafish larvae at 144 hpi. *Tg(mpeg1::Gal4-FF)*^gl25^/*Tg(UAS:LIFEACT-GFP)*^mu271^ larvae (with macrophages in green) were injected at 3 dpf in the hindbrain ventricle with 1000 CFU mCherry-labelled *S. sonnei* 53G (red). An individual infected macrophage harbouring persistent bacteria is magnified. Scalebars: 100 μm (left); 10 μm (right, inset). (**B**) 40% of bacterial fluorescence colocalises with *mpeg1*^*+*^ macrophages at persistent infection stages. (**C**) Individual macrophage harbouring mCherry-*S. sonnei*, followed over time for 12 h (from 24 to 36 hpi). For data in (C), 1000 CFU bacteria were delivered systemically in 2 dpf larvae. Scalebar: 10 μm. (**D**) *S. sonnei* 53G induces a reduced level of caspase-1 activation *in vivo* in the zebrafish model, as compared to *S. flexneri* M90T. For data in D, 10000 CFU bacteria were delivered systemically in 3 dpf larvae. Statistics: one-way ANOVA with Sidak’s correction; ***p<0.001; ****p<0.0001.

Previous work indicated that *S. sonnei* may be less efficient in inducing macrophage death than *S. flexneri* M90T (5a) *in vitro* using THP1 cells^27,29^. We hypothesised that this reduced ability to induce macrophage death may explain the ability to establish persistent infection of macrophages. To test this *in vivo*, we infected zebrafish larvae systemically (an infection route that promotes *Shigella*-macrophage interactions^29,30^) with *S. sonnei* 53G or *S. flexneri* 5a M90T. At 4 hpi, we dissociated infected larvae and PBS injection control larvae and quantified the level of caspase-1 activity (a marker of macrophage pyroptotic cell death) by flow cytometry (**Figure 3D**). Infection with *S. flexneri* 5a M90T led to a significantly higher level of caspase-1 activity compared to infection with *S. sonnei* 53G, supporting the conclusion that persistent *Shigella* serotypes fail to induce macrophage death *in vivo*.

## Conclusion

The zebrafish infection model has been instrumental to study host-pathogen interactions in response to a wide variety of human pathogens, including different *Shigella* species^31–34^. Here, we show that different *Shigella* serotypes (*S. sonnei, S. flexneri* 2a and *S. flexneri* 3a) that are highly prevalent in the MSM community and associated with direct host-to-host transmission, also establish persistent infection in zebrafish. Our results show that O-Ag variants promote establishment of persistent infection, but it is not yet known how the O-Ag precisely contributes. *S. sonnei* O-Ag is unique among enterobacterial pathogens and does not share obvious structural homology with the O-Ag expressed by *S. flexneri* 2a or 3a. However, *Shigella* species are a remarkable example of convergent evolution^23^ and the persistence phenotype mediated by different O-Ag serotypes may represent an example of pathoadaptive convergence due to occupation of the same niche. During the persistence phase, bacteria frequently co-localise with macrophages *in vivo*, strongly suggesting that macrophages represent a preferred niche for *Shigella* persistence. Strikingly, *S. sonnei* (which is highly associated with persistence) induces significantly less macrophage death *in vivo*, as compared to the *S. flexneri* serotype 5a (which has not been linked to persistence and MSM transmission). Considering that persistent infection of zebrafish recapitulates the epidemiological trend of persistence in humans, we propose that zebrafish can be used to discover host and pathogen factors underlying *Shigella* persistent infection in humans.

## STAR Methods RESOURCE AVAILABILITY

### Lead contact

Further information and requests for resources and reagents should contact Serge Mostowy.

### Materials availability

*Shigella* GM strains generated in this study are available upon request.

### Data and code availability

Sequencing data generated in this study have been deposited in the Sequencing Read Archive at BioProject number <u>PRJNA869897</u> and are publicly available as of the date of publication. Accession numbers are listed in the key resources table and against individual strains in **Table S1**.

## EXPERIMENTAL MODEL AND SUBJECT DETAIL

### Zebrafish model

Animal experiments were approved by the Home Office (project licenses Project license: PPL P4E664E3C) and performed following the Animals (Scientific Procedures) Act 1986. Wild-type AB zebrafish were used for all survival assays and CFU quantification experiments. For experiments involving imaging of zebrafish macrophages, these cells were labelled by crossing lines carrying the following the *Tg(mpeg1::Gal4-FF)*^gl25^ and *Tg(UAS:LIFEACT-GFP)*^mu271^ alleles. Eggs were obtained by naturally spawning, and both control and infected larvae were maintained at 28.5°C in embryo medium (0.5x E2 medium supplemented with 0.5 parts per million (ppm) of methylene blue. For injections and live microscopy, larvae were anaesthetized with 200 µg/ml tricaine (Sigma-Aldrich) in embryo medium. Injections were performed in larvae at 2 or 3 days post-fertilisation (dpf) and infected larvae were monitored up to 6 days post-infection. Sex was not determined, as all experiments were concluded before the stage of zebrafish sexual development.

## METHODS DETAILS

### Bacterial strains

Wild-type and genetically modified bacterial strains used in this work are further detailed in **Table S1 and Table S2**, respectively. Parental strains were made fluorescent by electroporation of pFPV25.1 (GFP labelled) or pFPV-mCherry (mCherry labelled), conferring Carbenicillin resistance. Exception was made for *S. sonnei* 373976, which was intrinsically resistant to high concentrations of Carbenicillin (MIC> 8,000 μg/ml) and could not be transformed. Bacterial mutant strains were previously described in Torraca et al., 2019^33^.

### Zebrafish infection

Bacteria were cultured at 37°C overnight in Trypticase Soy Broth (TSB) supplemented (when appropriate) with 100 ug/ml Carbenicillin, diluted 50x in fresh medium, and grown to Log phase. For inoculum preparation, bacteria were then spun down, washed in PBS and resuspended to an OD_600_ = 2 in an injection buffer containing 2% polyvinylpyrrolidone (Sigma-Aldrich) and 0.5% phenol red (Sigma-Aldrich) in PBS. Unless otherwise specified in the figure legend, 1 nl (corresponding to approximately 1000 CFU) of bacterial suspension was microinjected in the hindbrain ventricle (HBV) of 3 dpf zebrafish larvae. Exceptions include data in **Figure 3C, Figure S3C, Video S2** and **Figure 3D**. In these cases, 1000 CFU (**Figure 2C, Figure S3C, Video S2**) were delivered systemically in 2 dpf larvae or 10000 CFU (**Figure 3D**) were delivered systemically in 3 dpf larvae via the caudal vein.

### Survival assays and bacterial burden

Larvae failing to produce a heartbeat during a 30-second observation period were considered nonviable. For larvae beyond 5 dpf, clinical scoring criteria were applied to identify humane endpoints. Briefly, larvae were discontinued when the bacterial load (as assessed by live microscopy) was predictive of death in the following 24 hours (i.e., the hindbrain was filled with bacteria and/or infection was disseminated throughout the body) or when larvae resulted irresponsive to touch. These larvae were withdrawn from the experiment, euthanised and considered to be nonviable at the following timepoint.

For enumeration of live bacteria by CFU plating, larvae were washed in PBS, anesthetised, and mechanically homogenised in 200 μl (non-persistent stages, i.e., < 72 hpi) or 40 μl (persistent stages, i.e., ≥72 hpi) of PBS. Homogenates were serially diluted and plated onto Trypticase Soy Agar (TSA) containing 0.01% Congo red. Only viable larvae were used for CFU analysis.

### Clonality assay

Larvae were infected as previously specified, but with an inoculum composed of GFP-labelled *S. sonnei* and mCherry-labelled *S. sonnei* in a 1:1 ratio. The ratio between the two strains was determined by CFU plating at 0 hpi and 144 hpi. At 0 hpi, appropriate dilutions of the samples were made before plating, to correct for any differences in the occurrence of stochastic bottlenecks, due to different sample sizes.

### Microscopy

For *in vivo* time-lapse imaging, larvae were immobilised in 1% low-melting-point agarose. For high-resolution confocal microscopy, larvae were positioned in 35-mm-diameter glass-bottom MatTek dishes and imaging was performed using a Zeiss LSM 880 with fast Airyscan and 20× or 40× water immersion objectives. Image files were processed using ImageJ/FIJI software.

### Drug treatments

Nalidixic acid 1 μg/ml was directly added to the embryo medium. In pilot experiments, a range of concentrations were tested (0.5-4 μg/ml) and 1 μg/ml was the lowest concentration tested that fully prevented bacterial growth and death of zebrafish larvae upon infection with a lethal dose of *S. sonnei* 53G.

### Flow cytometry and Caspase-1 activity quantification

A pool of 3 dpf *Tg(mpeg1::Gal4-FF)*^gl25^/*Tg(UAS:LIFEACT-GFP)*^mu271^ larvae were infected systemically with 10000 CFU of mCherry-labelled *S. sonnei* 53G, *S. flexneri* M90T or PBS containing mock solution. At 4 hpi larvae were dissociated by treatment with 1 ml of 4% trypsin for 15 minutes at 28ºC. Single-cell dissociation was facilitated by mechanical disruption using a P1000 pipette after trypsin treatment. Dissociated cells were harvested by centrifugation (5 min, 800xg at room temperature), washed in calcium-free PBS, separated by passage on a 4 μm cell strainer and suspended in 500 μl staining solution for active caspase-1 (FLICA^®^ 660 Caspase-1 assay, probe: 660-YVAD-FMK, Immunochemistry technologies, #9122) prepared as per the manufacturer’s guidelines. Upon staining, cells were pelleted, washed in PBS and finally fixed in 4% paraformaldehyde overnight. For flow cytometry, cells were washed with 2 ml PBS twice and resuspended in 300 μl PBS. Active caspase-1 staining of single macrophages was measured on a LSRII (BD Biosciences) and data was analysed with the software FlowJo version 10.7.1.

### DNA extraction and sequencing

Genomic DNA was extracted from overnight bacterial cultures using the MasterPure Complete DNA Purification Kit (Lucigen Corporations, Wisconsin, US) according to manufacturer’s instructions. DNA concentration, purity and quantity was assessed using Nanodrop spectrophotometry (DeNovix, Thermofisher, UK) and Qubit fluorometer (Invitrogen, US) according to manufacturers’ instructions. Sequencing libraries were prepared using the ligation sequencing kit-SQK-LSK109 kit according to manufacturer’s instructions (Oxford Nanopore Technologies, Ltd, Oxford, UK) with slight modifications to the input DNA amounts (i.e., the input DNA amounts were increased at least two-fold at the initial step). DNA libraries were sequenced using a MinION sequencer and FLO-MIN106 Flow Cell version R9.4.1 (Oxford Nanopore Technologies Ltd, Oxford, UK).

### Assembly and annotation

The fast5 read files generated from the MinION instrument were base-called and demultiplexed with Guppy v5.0 (Oxford Nanopore Technologies Ltd. Oxford, UK). Processed read files were filtered using Filtlong v0.2.0, then assembled using Flye v2.9^35^. Racon v1.5.0^36^ was used to polish contigs with the nanopore reads and Medaka v1.6.1 (https://github.com/nanoporetech/medaka) was used to polish Racon polished contigs, with nanopore reads specifying the model r941_min_sup_g507. Polypolish v0.5.0 was then used to polish with Illumina reads, where they were available^37^. The quality and statistics of each assembly was evaluated with QUAST v4.4.0 without a reference genome^38^. Genomes were annotated using the Prokaryotic Genome Annotation Pipeline (PGAP). The complete genome sequence data have been submitted to the National Centre for Biotechnology (NCBI) and have been deposited at GenBank under the BioProject number <u>PRJNA869897</u>.

### Phylogenetic tree construction

Annotations of new genomes for the tree construction were performed using the Genome Annotation service made available by the Bacterial and Viral Bioinformatics Resource Centre (BV-BRC, https://www.bv-brc.org/)^19^. For the genome annotation, parameters were set as Annotation recipe: Bacteria/Archaea; Taxonomy name: *Shigella*. Phylogenetic trees were constructed using the Codon Tree method via the Phylogenetic tree building service made available by BV-BRC. For the tree building, parameters were set as Number of genes: 1000; Max allowed deletions: 1; Max allowed duplications: 1. Trees were constructed separately for *S. sonnei* and *S. flexneri* isolates. Data included the complete genomes of *S. sonnei* and *S. flexneri* isolates of known serotypes (publicly available via the BV-BRC database as for November 2022), as well as the draft genomes of all the isolates sequenced/resequenced and tested in this study (and reported in **Table S1**). The trees also include the draft genomes for isolates that had previously been reported to represent likely cases of carriage in humans with a distance between the collected isolates of at least 10 days^18^. Since this dataset represented pairs of isolates with <8 SNP distance, only sequences from the second isolates were used for tree construction. Data were visualised and annotated using FigTree v1.4.4. (http://tree.bio.ed.ac.uk/software/figtree/) and microreact (https://microreact.org/).

## QUANTIFICATION AND STATISTICAL ANALYSIS

Statistical tests were performed using Prism software (GraphPad Software, Inc.) or Microsoft Excel. The statistical significance of survival curves was determined using the log-rank Mantel-Cox test. In all other cases, statistical significance was determined using an unpaired two-tailed t-test or one-way ANOVA with Sidak correction, as further specified in the figure legends. Analyses were performed on Log10-transformed valued for CFU counts. Data are represented as mean ± standard errors of the mean (SEM).

## KEY RESOURCES TABLE

## Acknowledgements

We thank the Wiebke Herzog lab for the Tg(UAS:LIFEACT-GFP)mu271 zebrafish line and all members of the Mostowy, Baker and Holt labs for discussion and experimental help. V.T. was supported by an LSHTM/Wellcome Institutional Strategic Support Fund (ISSF) Fellowship (204928/Z/16/Z). D.B. is supported by the Deutsche Forschungsgemeinschaft (DFG) Walter Benjamin Programme (BR 6637/1-1). Research in K.B. laboratory is supported by the Academy of Medical Sciences Springboard award (SBF002\1114), a Medical Research Council New Investigator award (MR/R020787/1), and the BBSRC (BB/V009184/1). Work in the K.E.H laboratory is supported by the Bill and Melinda Gates Foundation, Seattle (OPP1175797), KlebNet Project. Research in S.M. laboratory is supported by a European Research Council Consolidator Grant (772853 - ENTRAPMENT), Wellcome Trust Senior Research Fellowship (206444/Z/17/Z) and the Lister Institute of Preventive Medicine. K.B. and C.J. are affiliated with the National Institute for Health Research Health Protection Research Unit (NIHR HPRU) in Gastrointestinal Infections at the University of Liverpool in partnership with the United Kingdom Health Security Agency (UKHSA), in collaboration with the University of Warwick. The views expressed are those of the author(s) and not necessarily those of the NHS, the NIHR, the Department of Health and Social Care or UKHSA.

## Author contributions

This project was conceived by V.T. and S.M. V.T. performed all zebrafish experiments and analysed all datasets. V.T., S.L.M., C.E.C. and M.D.S. prepared DNA samples and analysed sequencing results. V.T. constructed the phylogenetic trees with inputs from K.E.H. and K.B. V.T and D.B. performed experiments to detect active caspase-1 in zebrafish. S.B. and C.J. provided valuable strains for the work. V.T. and S.M. wrote the manuscript with input from all the authors.

## Supplementary Figure legends

**Figure S1 (related to Fig. 1). *Shigella* establishes persistent infection in zebrafish** (**A-C**) Linear regression analysis of the change of Log_10_(CFU) of *S. sonnei* as a function of time for untreated larvae (A) or larvae treated with the antibiotic Nalidixic Acid (B), and relevant summary statistics (C). In the absence of antibiotics, *Shigella* infection progresses in three distinct phases (acute, clearing and persistent), that can be described by individual linear equations. Treatment with therapeutic doses of Nalidixic acid prevents infection from undergoing acute expansion, but it does not promote complete eradication of the infection. The presence or absence of the antibiotic does not significantly impact the trend (slope) of Log_10_(CFU) over time, neither during the clearing phase nor during the persistence phase. (**D-F**) Survival curves (D), Log_10_(CFU) (E) and phenotypic analysis (F) of zebrafish larvae infected with a lethal dose (i.e. 8000 CFU) of *S. sonnei* and treated with different concentrations of Nalidixic acid. 1 μg/ml Nalidixic acid was sufficient to completely rescue host survival (D) and prevent significant bacterial growth *in vivo* during the acute infection phase (E). Toxicity at this antibiotic concentration is negligible (F) and <10% of treated larvae displayed aberrant phenotypes (i.e., cardiac oedemas). (**G**) *Shigella* persistent infection is associated with antibiotic tolerance. At 72 hpi (i.e., end of the clearing phase) only 0.53% of the tested larvae carried bacteria that developed an inheritable decreased susceptibility to the antibiotic treatment (i.e. 4x MIC).

**Figure S2 (related to Fig. 2). *Shigella* O-Antigen serotype associated with MSM transmission enables persistent infection**

(**A-B**) Log_10_CFU of individual larvae (A) and survival curves (B), for WT *S. sonnei* 53G and several isogenic mutants. (**C-D**) Log_10_CFU of individual larvae (C) and survival curves (D) for several *S. sonnei* isolates from Lineage II and Lineage III. (**E-F**) Log_10_CFU of individual larvae (E) and survival curves (F) for several *S. flexneri* isolates from serotypes 2a, 3a and 5a. (**G-H**) Phylogenetic trees (codon trees) for *S. sonnei* (G) and *S. flexneri* (H) constructed by using complete reference genomes of known serotypes (available in BV-BRC, https://www.bv-brc.org/), draft genome sequences of isolates associated with persistence carriage in humans (available from Bengtsson et al.^18^) and the isolates used in this study (sequenced with nanopore technology). Red labels indicate strains associated with persistent carriage in humans, orange labels indicate strains that established persistent infection in zebrafish, while black labels indicate strains that did not establish persistent infection in zebrafish. For the *S. sonnei* tree, different lineages (II and III) and sub-lineages (3.6. and 3.7) are highlighted with different colours. For the *S. flexneri* tree, the three main clusters encompassing the serotypes investigated in this study (2a, 3, and 5) are highlighted with different colours and the serotype of all the strains is reported in brackets after the strain name. Statistics: one-way ANOVA with Sidak’s correction on Log10-transformed data (A, C, E); Log-rank Mantel-Cox test (B, D, F); ns (non-significant) p≥0.05; *p<0.05; **p<0.01; ***p<0.001; ****p<0.0001.

**Figure S3 (related to Fig. 3). *Shigella* can establish persistent infection of macrophages *in vivo***

(**A**) Head region and individual macrophage detail from a representative *S. sonnei* infected zebrafish larvae at 72 hpi. *Tg(mpeg1::Gal4-FF)*^gl25^/*Tg(UAS:LIFEACT-GFP)*^mu271^ larvae (with macrophages in green) were injected at 3 dpf in the hindbrain ventricle with 1000 CFU mCherry-labelled *S. sonnei* 53G (red). A cluster of infected macrophages harbouring persistent bacteria is magnified. Scalebars: 100 μm (left); 10 μm (right, inset). (**B**) Longitudinal imaging of the head region from a representative *S. sonnei* infected zebrafish larvae at 0, 24, 72 and 144 hpi. The same region of interest is magnified in the bottom left corner. Scalebar: 100 μm. (**C**) Longitudinal quantification of bacterial fluorescence in an individual macrophage harbouring mCherry-*S. sonnei*, followed over time for 12 h (from 24 to 36 hpi). The bacterial fluorescence remains overall constant or only slightly increases (i.e., bacterial fluorescence shows only a 1.21 ± 0.21 fold change over a 12 h period of observation). For data in C, 1000 CFU bacteria were delivered systemically in 2 dpf larvae. Scalebar: 10 μm.

**Table S1. Bacterial strains used in this study**

**Table S2. Details of newly sequenced/re-sequenced bacterial strains used in this study**

**Video S1 (related to Figure 3). Confocal z-stack of a macrophage carrying *S. sonnei* infection at 144 hpi**.

*Tg(mpeg1::Gal4-FF)*^gl25^/*Tg(UAS:LIFEACT-GFP)*^mu271^ larvae (with macrophages labelled in green) were injected at 3 dpf in the hindbrain ventricle with 1000 CFU mCherry-labelled *S. sonnei* 53G (red). Images were taken by confocal microscopy at 144 hpi.

**Video S2 (related to Figure 3). Confocal time lapse of a macrophage carrying *S. sonnei* infection for 12 h**.

*Tg(mpeg1::Gal4-FF)*^gl25^/*Tg(UAS:LIFEACT-GFP)*^mu271^ larvae (with macrophages labelled in green) were injected at 2 dpf via the caudal vein with 1000 CFU mCherry-labelled *S. sonnei* 53G (red). Images were taken by confocal microscopy at 6.5 minute intervals for 12 h, from 24 to 36 hpi.

