## Supplementary material for "*Shigella* serotypes associated with carriage in humans establish persistent infection in zebrafish": Figure S1

### Figure S1 (related to Fig. 1). *Shigella* establishes persistent infection in zebrafish

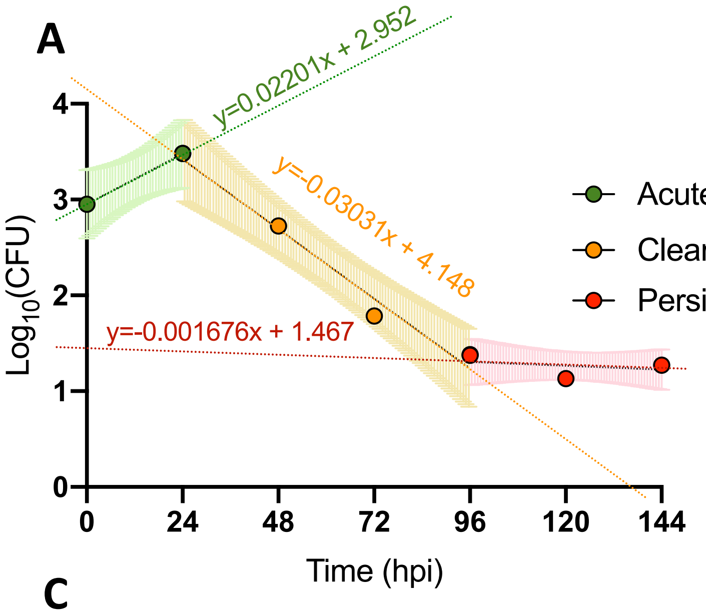


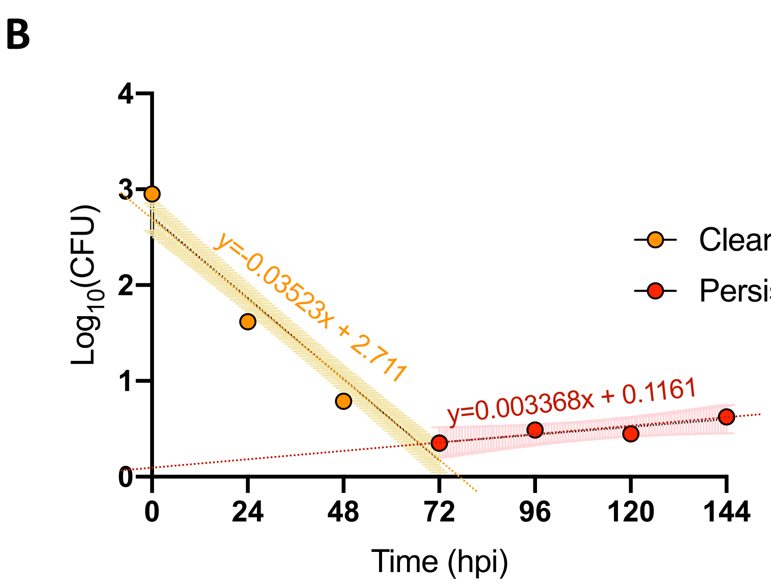


**C**

| | Untreated Acute | Untreated Clearing | Untreated Persistent | 1 µg/ml Nal Clearing | 1 µg/ml Nal Persistent |
|---|---|---|---|---|---|
| Untreated Clearing | p=0.0005 | - | - | - | - |
| Untreated Persistent | p=0.0200 | p<0.0001 | - | - | - |
| 1 µg/ml Nal Clearing | p<0.0001 | ns p=0.3165 | p<0.0001 | - | - |
| 1 µg/ml Nal Persistent | p=0.0204 | p<0.0001 | ns p=0.1750 | p<0.0001 | - |

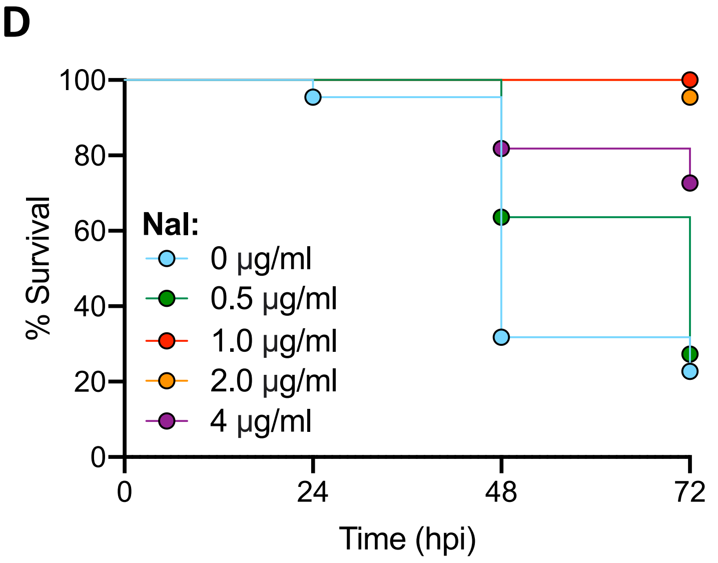


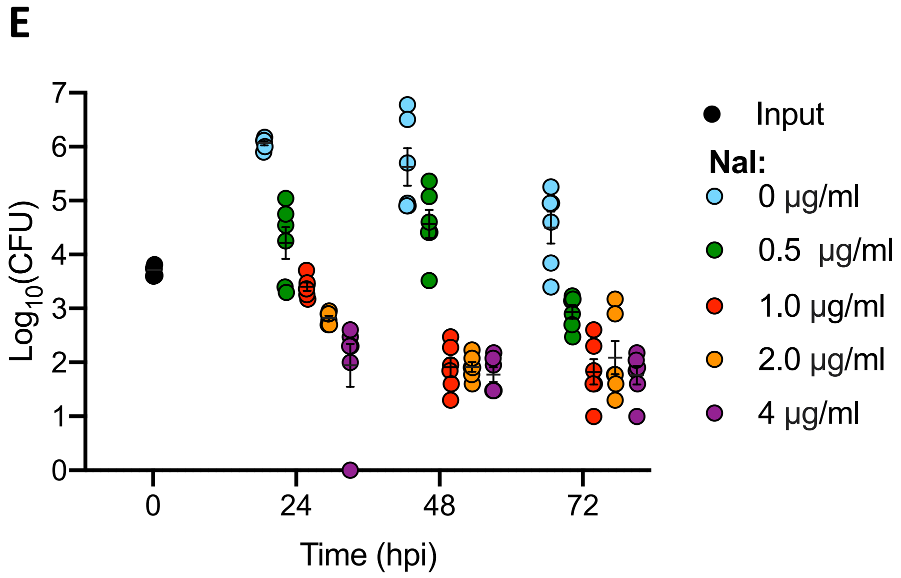


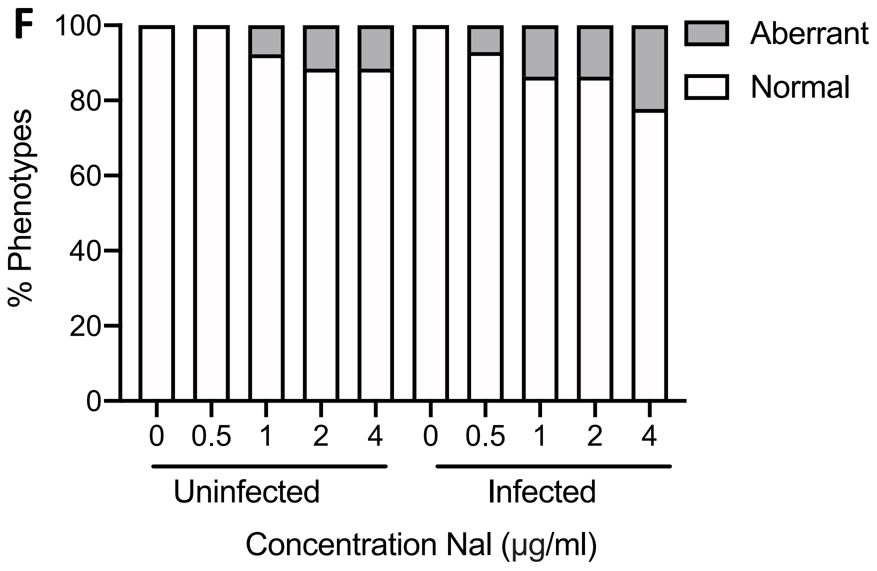


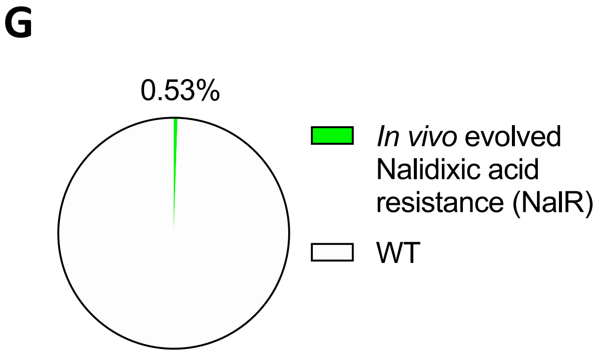
