## Supplementary material for "*Shigella* serotypes associated with carriage in humans establish persistent infection in zebrafish": Figure S2

**Figure S2 (related to Fig. 2). *Shigella* O-Antigen serotypes associated with MSM transmission enable persistent infection**

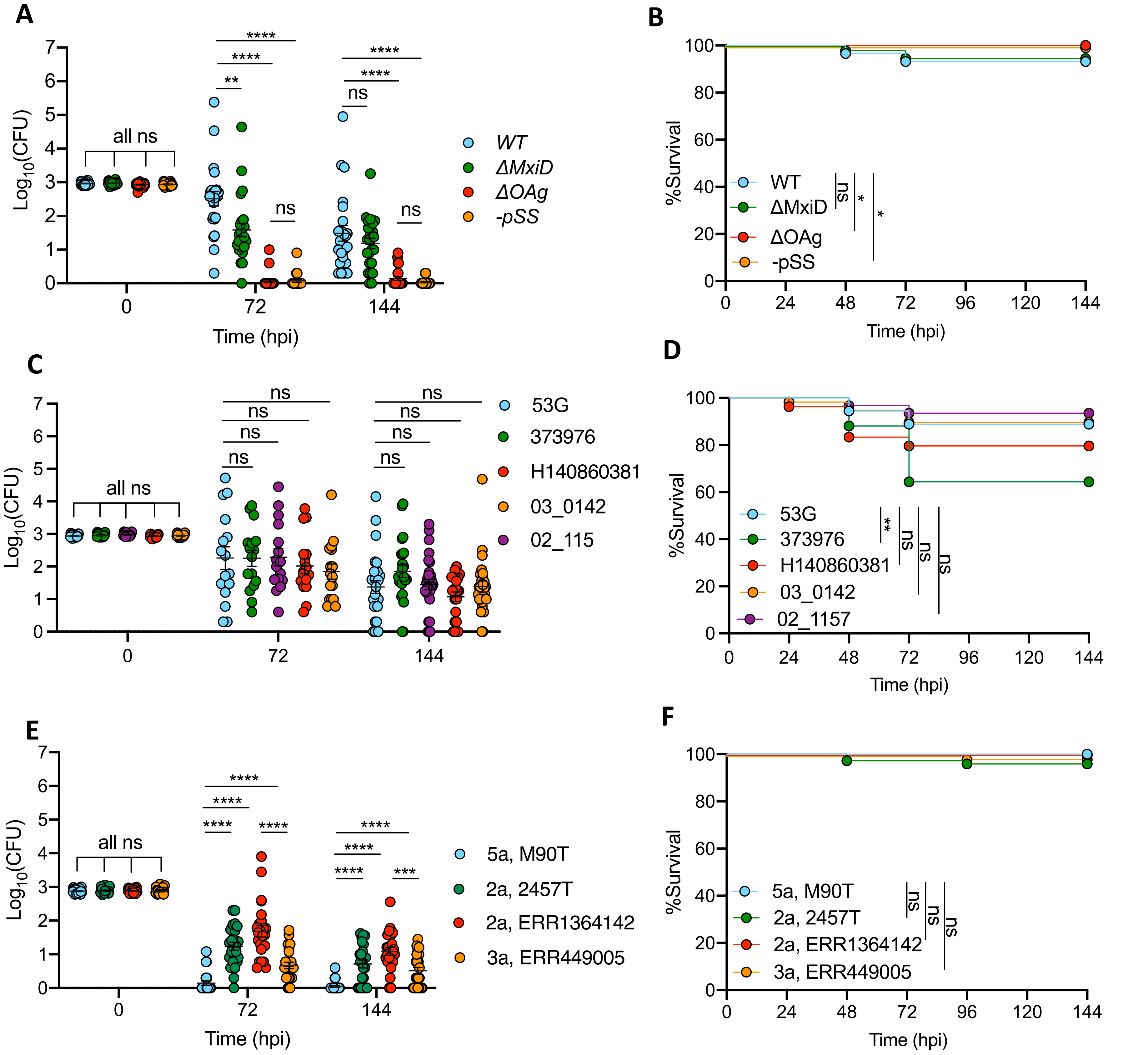

**Figure S2 - continued. Supplement to Fig. 3. *Shigella* O-Antigen serotypes associated with MSM transmission enable persistent infection**

G

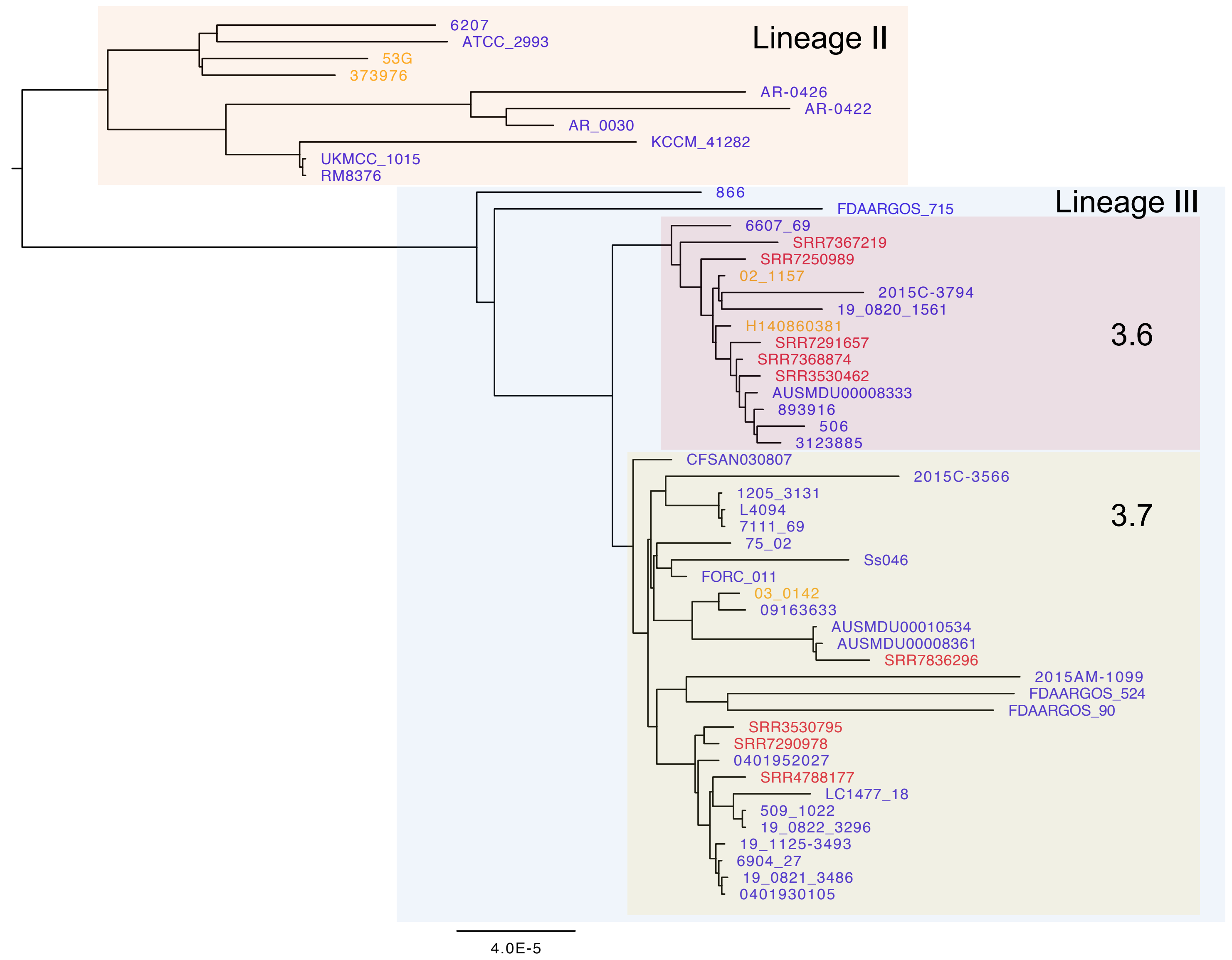


H

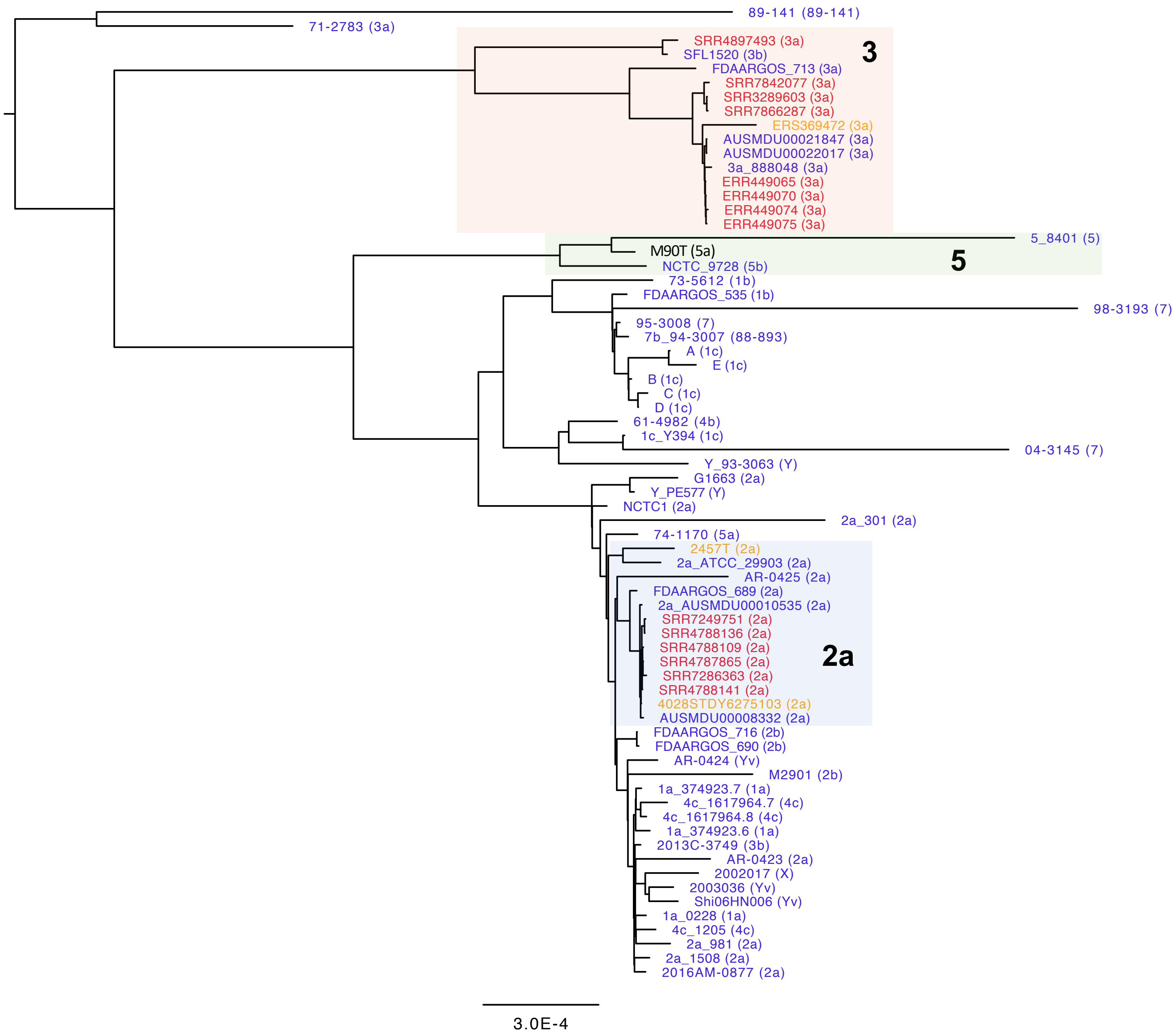
