## Supplementary figures and images for "*Shigella* serotypes associated with carriage in humans establish persistent infection in zebrafish"

### Figure S3

**Figure S3 (related to Fig. 3). *Shigella* can establish persistent infection of macrophages *in vivo***

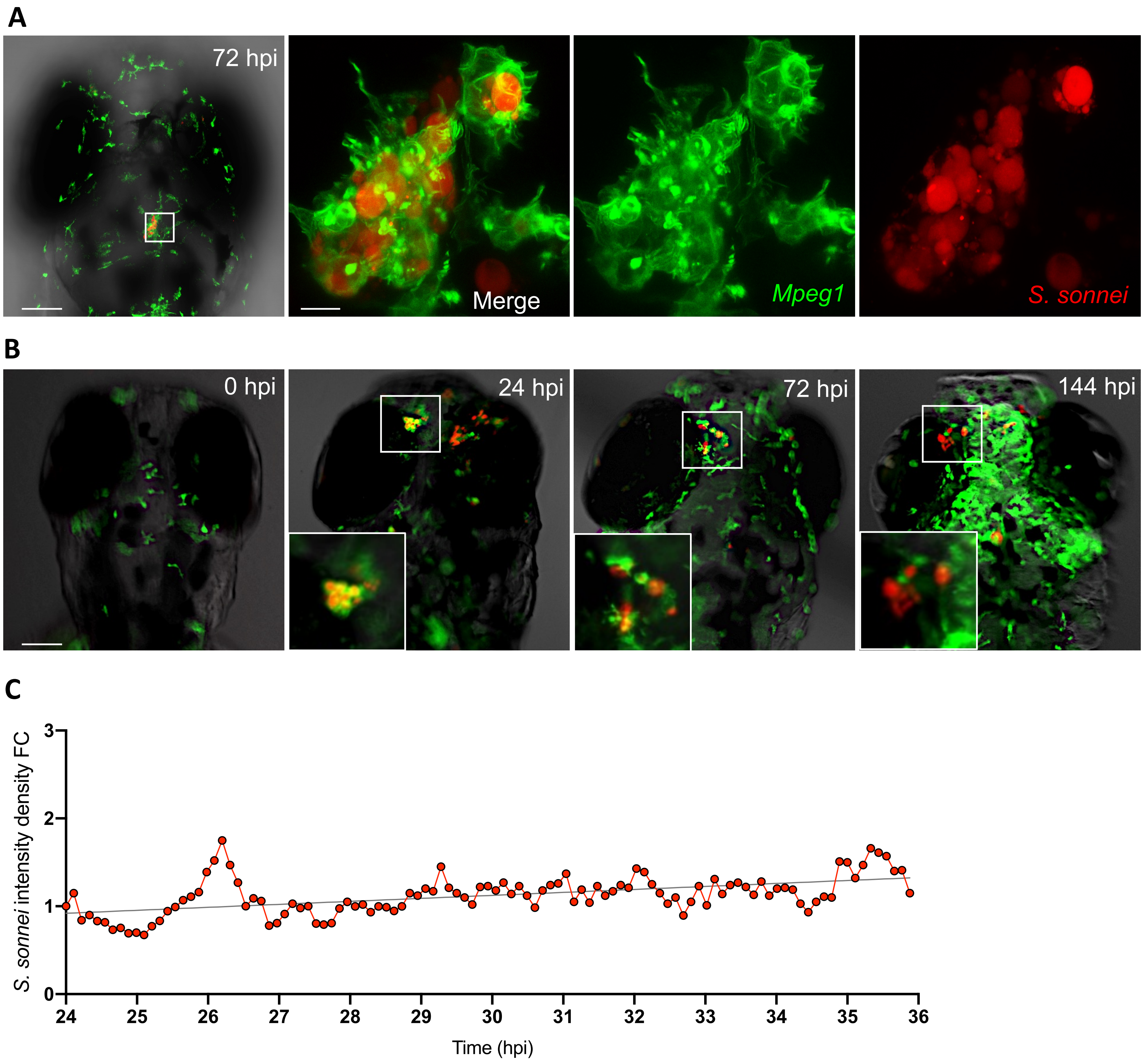
