## Supplementary material for "*Shigella* serotypes associated with carriage in humans establish persistent infection in zebrafish": Table S1

**Table S1. Details of newly sequenced/re-sequenced bacterial strains used in this study**

| **Species** | **Serotype** | **Strain Name** | **Lineage (Genotype)** | **Reference** | **Notes** | **Sequence accession** |
| --- | --- | --- | --- | --- | --- | --- |
| *Shigella sonnei* | Ss | 53G | II (2.8) | Formal et al., 1966 | 1954 isolate from Japan, widely used in the lab | [GCF_000283715.1](https://www.ncbi.nlm.nih.gov/assembly/GCF_000283715.1/) |
| *Shigella sonnei* | Ss | 373976 | II (2.12.4) | Bardsley et al., 2020 | 2017 isolate from the UK (UKHSA , C. Jenkins) | [GCA_026013135.1](https://www.ebi.ac.uk/ena/browser/view/GCA_026013135.1) |
| *Shigella sonnei* | Ss | H140860381 | III (3.6.1.1) | Baker et al., 2016 | 2017 isolate from the UK (UKHSA , C. Jenkins) | [ERR861624](https://www.ebi.ac.uk/ena/browser/view/ERR861624) |
| *Shigella sonnei* | Ss | 03_0142 | III Global (3.7.29.1.4) | The et al, 2019 | 2017 isolate from the UK (UKHSA , C. Jenkins) | [GCA_026013065.1](https://www.ebi.ac.uk/ena/browser/view/GCA_026013065.1) |
| *Shigella sonnei* | Ss | 02_1157 | III CenAsia (3.6.1.1) | The et al, 2019 | 2014 isolate from Bhutan (S. Baker) | [GCA_026013055.1](https://www.ebi.ac.uk/ena/browser/view/GCA_026013055.1) |
| *Shigella flexneri* | 5a | M90T | - | Sansonetti et al., 1982 | 1981 isolate from the United States, widely used in the lab | [GCA_026013175.1](https://www.ebi.ac.uk/ena/browser/view/GCA_026013175.1) |
| *Shigella flexneri* | 2a | 2457T | - | Formal et al., 1958 | 1954 isolate from Japan, widely used in the lab | [GCF_000007405.1](https://www.ncbi.nlm.nih.gov/assembly/GCF_000007405.1) |
| *Shigella flexneri* | 2a | 4028STDY6275103 | - | Baker et al., 2018 | 2017 isolate from the UK (UKHSA , C. Jenkins) | [GCA_026013095.1](https://www.ebi.ac.uk/ena/browser/view/GCA_026013095.1) |
| *Shigella flexneri* | 3a | ERS369472 | - | Allen et al., 2021 | 2011 isolate from the UK (UKHSA , C. Jenkins) | [GCA_026013105.1](https://www.ebi.ac.uk/ena/browser/view/GCA_026013105.1) |
