## Supplementary material for "*Shigella* serotypes associated with carriage in humans establish persistent infection in zebrafish": Table S2

**Table S2. Genetically modified bacterial strains used in this study**

| **Species** | **Serotype** | **Strain background** | **Reporter construct** | **Mutations** | **Reference** |
| --- | --- | --- | --- | --- | --- |
| *Shigella sonnei* | Ss | 53G | GFP from pFPV25.1 (Valdivia et al., 2006) | WT | Torraca et al., 2019 |
| *Shigella sonnei* | Ss | 53G | GFP from pFPV25.1 (Valdivia et al., 2006) | ΔMxiD (Watson et al., 2018) | Torraca et al., 2019 |
| *Shigella sonnei* | Ss | 53G | GFP from pFPV25.1 (Valdivia et al., 2006) | ΔOAg (Watson et al., 2019) | Torraca et al., 2019 |
| *Shigella sonnei* | Ss | 53G | GFP from pFPV25.1 (Valdivia et al., 2006) | -pSS (Torraca et al., 2019) | Torraca et al., 2019 |
| *Shigella sonnei* | Ss | 53G | mCherry from pFPV-mcherry (Drecktrah et al., 2008) | WT | Torraca et al., 2019 |
| *Shigella sonnei* | Ss | H140860381 | mCherry from pFPV-mcherry (Drecktrah et al., 2008) | WT | This study |
| *Shigella sonnei* | Ss | 03_0142 | mCherry from pFPV-mcherry (Drecktrah et al., 2008) | WT | This study |
| *Shigella sonnei* | Ss | 02_1157 | mCherry from pFPV-mcherry (Drecktrah et al., 2008) | WT | This study |
| *Shigella flexneri* | 5a | M90T | mCherry from pFPV-mcherry (Drecktrah et al., 2008) | WT | This study |
| *Shigella flexneri* | 2a | 2457T | mCherry from pFPV-mcherry (Drecktrah et al., 2008) | WT | This study |
| *Shigella flexneri* | 2a | 4028STDY6275103 | mCherry from pFPV-mcherry (Drecktrah et al., 2008) | WT | This study |
| *Shigella flexneri* | 3a | ERS369472 | mCherry from pFPV-mcherry (Drecktrah et al., 2008) | WT | This study |
